# Physiological landmarks reveal the laminar organization of the primate hippocampus

**DOI:** 10.64898/2026.09.12.750217

**Authors:** Sofia M. Landi, Michael J. Jutras, Ellen C.S. Bakotich, Elizabeth A. Buffalo

**Affiliations:** Department of Neurobiology and Biophysics, University of Washington, Seattle, WA. USA; Washington National Biomedical Research Center, University of Washington, Seattle, WA. USA; Graduate Program in Neuroscience, University of Washington, Seattle, WA. USA

## Abstract

The laminar anatomy of the hippocampal microcircuit is conserved across mammals, and its physiology has been characterized extensively in rodents, yet how this circuitry is organized physiologically in the primate brain remains poorly understood. Here we used high-density Neuropixels probes to record simultaneously across the subfields of CA1, CA3 and the dentate gyrus in two awake macaques. Sharp-wave ripples and dentate spikes produced current source density signatures that provided physiological landmarks for CA1 and the dentate gyrus, enabling alignment of recordings to hippocampal cytoarchitecture. Theta-band (3-7 Hz) oscillatory activity occurred in discrete bouts whose prevalence peaked in the *stratum lacunosum-moleculare*. In CA1, gamma-band amplitude was modulated by theta-band phase, with fast and slow gamma preferentially coupled to distinct phases of the theta cycle. Putative principal cells and two interneuron classes differed systematically in firing rate, burstiness, and rhythmic modulation, with principal cells showing theta-rhythmic firing and wide-waveform interneurons showing alpha- and beta-rhythmic firing. Cross-correlogram analyses revealed sparse, distance-dependent functional coupling with a consistent directional hierarchy in which principal cells led interneurons. Together, these results identify the laminar microcircuit organization of the primate hippocampus *in vivo* and provide a new physiological framework for interpreting primate hippocampal recordings.

## Introduction

Hippocampal activity has been associated with memory formation and spatial processing. Neural computations that give rise to functions like these depend not only on which neurons are active, but also on how cells and synaptic pathways are organized within a circuit. The hippocampal formation provides a striking example of this spatial organization. Principal cell bodies are concentrated within distinct layers, while major intrinsic and extrinsic pathways terminate in spatially organized dendritic compartments across the dentate gyrus (DG), CA3, and CA1 (1–4). This laminar architecture provides an anatomical framework for interpreting hippocampal physiology because neural signals recorded at different depths can reflect different cell populations, synaptic inputs, and network processes. However, exploiting this framework in primates has been difficult. Although histological reconstruction can localize an electrode track post hoc (5), acute probes are repositioned across sessions, so histology provides a single snapshot at the end of an experiment rather than the per-session registration that interpreting depth-resolved signals requires. Compounding this, conventional probes have lacked the spatial resolution to sample densely across the full hippocampus. Recent laminar recordings in macaque CA1 have demonstrated that physiological signals associated with sharp-wave ripples (SWRs) can be used to align recordings across CA1 depth and resolve sublayer-specific organization (6). However, a physiological framework across the broader primate hippocampal formation has not been developed.

High-density, laminar recordings of local field potentials (LFPs) offer a means of establishing such a framework because the highly organized hippocampal anatomy gives rise to spatially localized extracellular current patterns. In rodents, current source density (CSD) analyses have identified laminar signatures associated with several hippocampal network events. SWRs are accompanied by a characteristic organization of transmembrane currents across CA1 dendritic and somatic layers, with ripple power peaking at the pyramidal cell layer (7). Dentate spikes (DSs) produce prominent sinks within the molecular layer of the dentate gyrus, reflecting activation of entorhinal inputs (8, 9). Theta-band activity likewise varies systematically across hippocampal depth in both amplitude and phase, with prominent theta-band oscillations near the hippocampal fissure and CA1 *stratum lacunosum-moleculare* (10, 11). Together, these signals provide candidate physiological fiducials for linking extracellular recordings to the underlying hippocampal cytoarchitecture. Importantly, hippocampal dynamics in primates cannot be simply inferred from rodent physiology: theta-band activity in primates is typically intermittent and differs in its behavioral expression and relationship to other rhythms (12–15), and hippocampal cell layer width and density differ across species (2). Resolving laminar organization in the primate therefore requires probes that combine dense laminar sampling with the single-unit yield and extent of anatomical coverage to capture the full functional diversity of hippocampal cell populations across layers.

Here, we used high-density Neuropixels recordings spanning CA1, CA3 and the dentate gyrus in awake macaques to establish a physiological framework for resolving hippocampal laminar organization *in vivo*. We used SWR- and DS-triggered CSD to identify complementary depth landmarks and test their convergence with the laminar distributions of neuronal firing and theta-band activity. This coordinate system allowed us to characterize the organization of theta and gamma oscillations and their cross-frequency coupling, the laminar distributions and rhythmic properties of putative neuronal classes, and fine-timescale functional interactions between cell types. Together, these analyses provide a comprehensive view of laminar microcircuitry in the primate hippocampus and a foundation for biologically grounded models of how this circuitry supports memory and cognition.

## Results

We recorded across hippocampal layers and subregions in two head-fixed macaques as they performed a virtual reality foraging task (5 sessions, monkey C) and a picture viewing task (7 sessions, monkey C and monkey M). Recordings were made with the probe oriented perpendicular to the horizontal stereotaxic plane, allowing simultaneous sampling across the full laminar depth of the hippocampal formation (Fig. 1A). Throughout, depth is reported relative to the dentate spike (DS) sink (0 µm; DS characterized below), with positive values towards CA3 and negative values towards CA1 (Fig. 1). Within each subfield, we use “superficial” and “deep” relative to the principal cell layer, with superficial denoting the dendritic and molecular layers that lie toward the hippocampal fissure. The density of recorded units of different cell types showed sharp transitions with depth, consistent with the dense packing of excitatory neurons into discrete layers (Fig. 1B).

**Figure 1:**
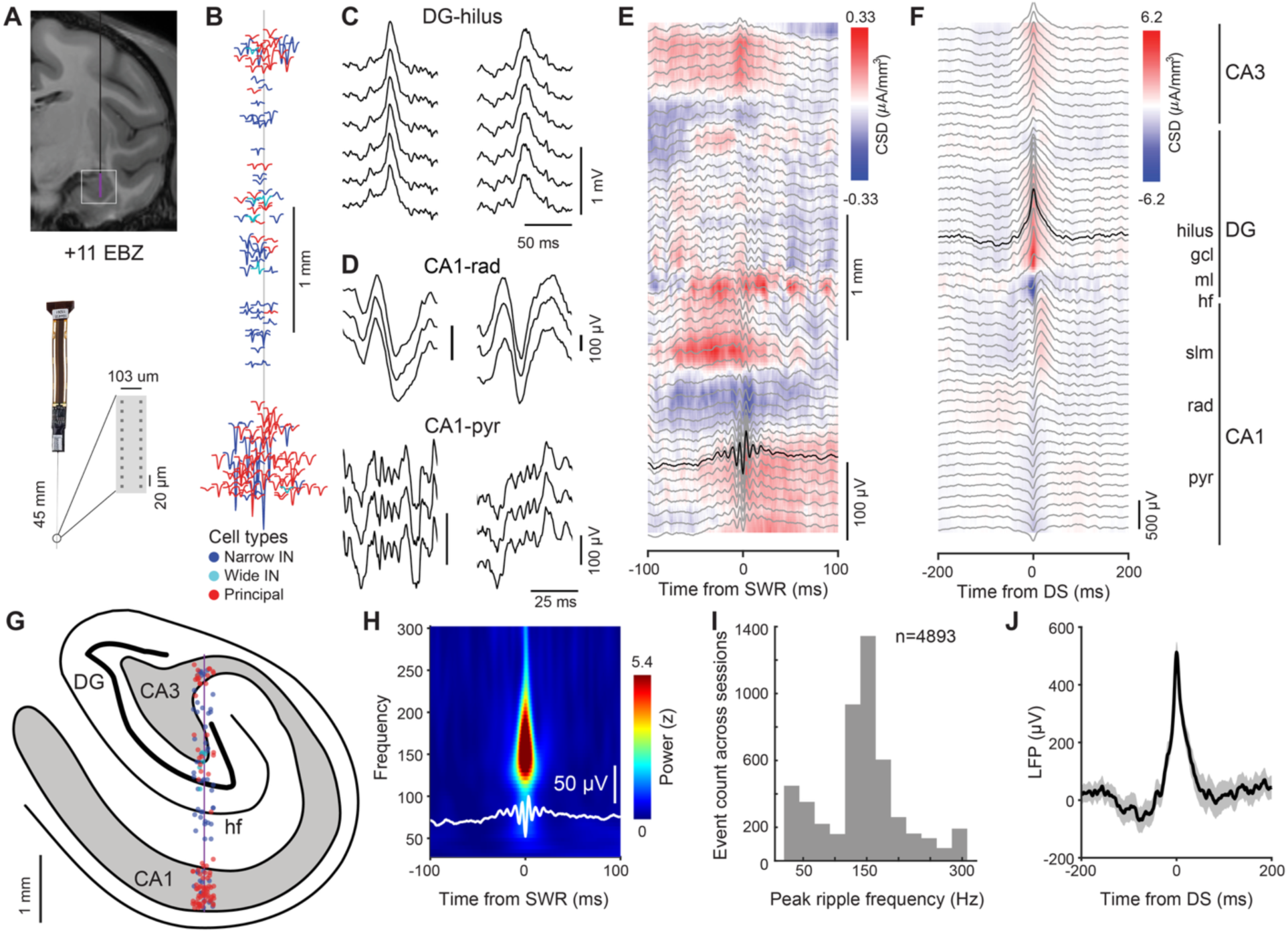
Physiological characterization of hippocampal layers and subregions. (**A**) Top: Magnetic resonance image (coronal) from Monkey C showing the recording hemisphere and a representative probe trajectory at 11mm anterior to ear bar zero (EBZ). Bottom: probe geometry and recording site layout. (**B**) Spatial distribution of putative single units in a representative recording session plotted across the probe depth at the channel of maximal spike waveform amplitude. Units are classified as principal cells (red), narrow-waveform interneurons (blue), and wide-waveform interneurons (cyan) based on spike waveform duration and autocorrelogram criteria. (**C**) Two sample dentate spikes across six channels adjacent to the maximum amplitude channel. (**D**) Two sample sharp-wave ripples, shown on three channels adjacent to the detection channel (putative CA1 pyramidal cell layer) and on three channels around the current sink superficial to the detection channel (putative CA1 stratum radiatum). (**E**) Average current source density (CSD) across all channels aligned to SWRs computed from n=283 events in one session, with the averaged LFP traces overlaid in gray, detection channel in black. (**F**) Average CSD across all channels aligned to DSs computed from n=28 events in one session, with the averaged LFP traces overlaid in gray, detection channel in black. (**G**) Estimate of probe shank position using DS, SWR and cell type profiles and landmarks, overlaid on a line drawing illustrating the cytoarchitectonic divisions of the hippocampus. (**H**) SWR-triggered time-frequency spectrogram computed by Morlet wavelet transform (30-300 Hz). Each event was z-scored per frequency to its own pre-event baseline, then averaged across events. White trace: mean ripple waveform. (**I**) Histogram of peak ripple frequency of all events (n=4893) detected in all 12 sessions. (**J**) Mean ±SEM of the LFP across recording sessions at the maximum amplitude DS channel aligned to the DS peak. Color bars indicate CSD amplitude (µA/mm³) in E and F and normalized power (z) in H. CA1, cornu ammonis 1; CA3, cornu ammonis 3; slm, stratum lacunosum-moleculare; rad, stratum radiatum; pyr, pyramidal cell layer (stratum pyramidale); hf, hippocampal fissure; DG, dentate gyrus; ml, molecular layer; gcl, granule cell layer; hilus, polymorphic layer.

In primates, the CA1 pyramidal layer is considerably thicker and more diffusely organized than in rodents (10-15 cells vs. ∼ 5 cells thick) (2, 4, 6, 16) and could therefore be identified as the thicker cell body layer enriched in putative excitatory neurons (Fig. 1B, Fig. S1). However, identifying DG and CA3 was less straightforward, and moreover, the extent of the dendritic compartments cannot be determined from unit density alone (3). We therefore asked whether network events could provide independent physiological landmarks for resolving the hippocampal layers *in vivo*. In rodents, sharp-wave ripples and dentate spikes are associated with stereotyped laminar current signatures, making them candidate signals for this characterization.

We identified dentate spikes (DS, Fig. 1C) and sharp-wave ripples (SWR, Fig. 1D) throughout each recording and constructed event-triggered average current-source density (CSD) maps to localize the underlying laminar current generators (Fig. 1E-F). These two physiologically-defined landmarks confirmed and refined the initial unit-density assignment and provided two independent physiological landmarks for cross-session alignment, each arising from a physically distinct current generator: the ripple-based reference for CA1 and the DS-based reference for the dentate gyrus, allowing us to align physiology to anatomy (Fig. 1G).

For each recording session, we identified the CA1 principal cell layer reference channel as the site with the highest ratio of ripple-band to broadband power, confirmed post hoc as the channel with the strongest mean ripple amplitude across detected events (Fig. 1D-E). This ripple-based reference confirmed the layer assignment obtained from unit density (Fig. 1B). Superficial to it, the CSD showed a characteristic sink-source reversal across the CA1 dendritic layers, which we assigned to a current sink in *stratum radiatum* and a current source in *stratum lacunosum-moleculare* (Fig. 1E), consistent with what has been described in rodents (7). Time-frequency decomposition of individual events confirmed the expected oscillatory signature in the ripple band (Fig. 1H) (17), with peak frequencies distributed around ∼150 Hz across all detected events (n = 4893, 12 sessions; Fig. 1I).

We applied a similar logic to identify the dentate gyrus, using DS-evoked CSD as a reference. In rodents, DSs are characterized by a current sink in the molecular layer of the dentate gyrus, reflecting excitatory input from the entorhinal cortex (8, 9), with a coupled, shallower source whose reversal marks the boundary of the granule cell layer (18). For each recording session, we selected the DS reference channel by ranking channels on DS-triggered amplitude and event count (Fig. 1C, 1F, 1J). Deep to this reference, the CSD reproduced the canonical sink-source arrangement (Fig. 1F), which we identified as the molecular and granule cell layers of the dentate gyrus. This provided a second depth reference, independent of the CA1 ripple landmark.

Event rates differed markedly between SWRs and DSs. Dentate spikes occurred at 1.43±0.10 events/min in the picture viewing task, and 1.61±0.27 events/min in the foraging task, whereas SWRs occurred at 13.46±1.28 and 14.83±0.46 events/min, respectively (mean ± SEM across sessions). Both event types showed a modest increase in the foraging task relative to the picture viewing task.

With the laminar framework in place, we next examined how oscillatory activity was distributed across hippocampal subregions and cell layers. Initial observations revealed that spectral power varied with depth in a frequency-specific manner, with different bands most prominent at different laminar positions. After removing the aperiodic (1/f) component of the power spectra using FOOOF (19), periodic peaks in the low frequency range were concentrated between 3-7 Hz, with notable variability across laminar contacts (Fig. 2A). We accordingly defined theta as 3-7 Hz for subsequent analyses. For each recording, we designated the contact with the highest FOOOF-corrected theta-band power as the theta reference channel, which was subsequently used as the common reference for the laminar theta analyses shown in Figures 2 and 3.

**Figure 2.**
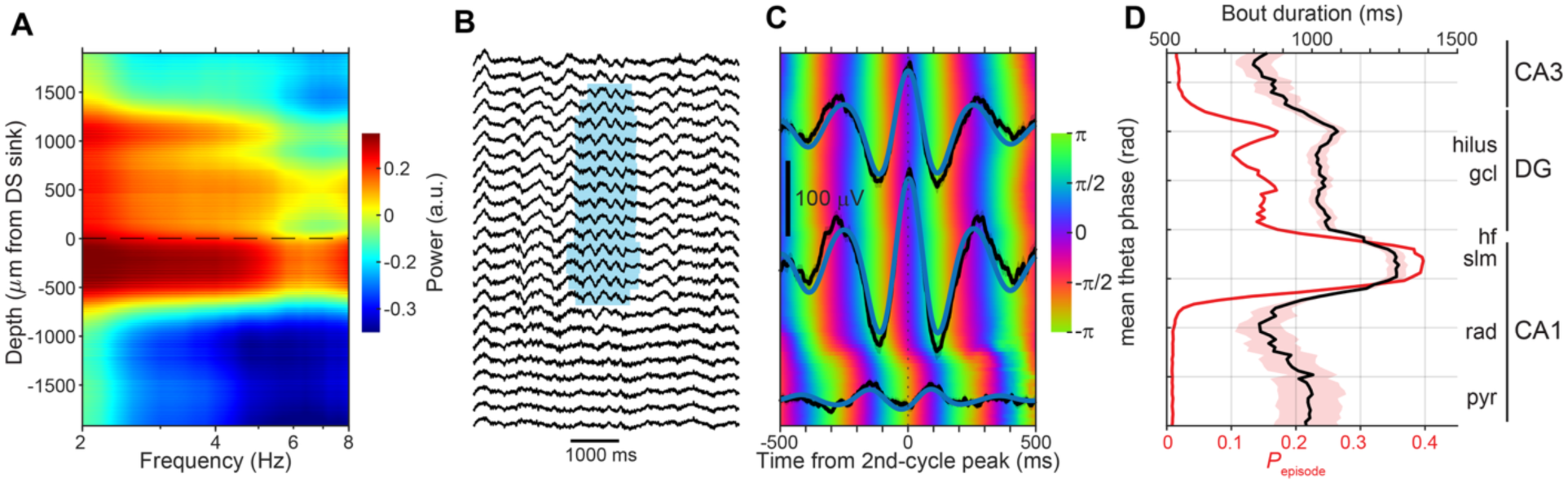
Theta-band oscillatory activity across hippocampal layers and subregions. (**A**) FOOOF-corrected power spectra at 2-8 Hz across all contacts spanning the hippocampus. (**B**) Representative theta-band oscillatory bout detected by *P*_episode_ highlighted in cyan. LFPs sampled from a subset of contacts across the probe. (**C**) Color plot portrays average circular mean of theta-band (3-7 Hz) phase across bouts aligned to the theta-band reference channel, peak-locked to the second cycle in the bout. Line plots represent average LFPs (black) and LFPs filtered in the theta band (blue) aligned to the same reference timepoint for three sample channels. (**D**) Theta-band *P*_episode_ values (red) and mean duration of theta bouts (black) for the sample recording; shading denotes SEM in the bout duration.

**Figure 3.**
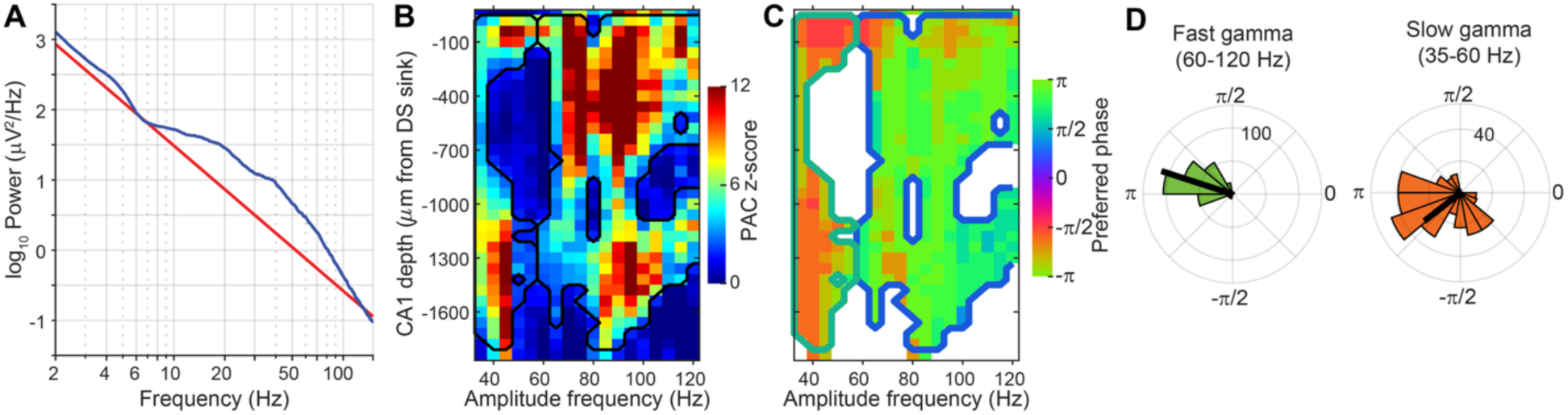
Gamma-band amplitude is modulated by theta-band phase. (**A**) Average power across CA1 contacts (blue) and FOOOF-derived aperiodic fit to median power spectrum (red) for a sample recording. (**B**) Phase-amplitude coupling between CSD theta-band phase (3-7 Hz) and LFP amplitude (35-120 Hz) across CA1 channels for a sample recording. Values are z-scored against 500 circular-shift surrogates; black contours show clusters from permutation testing at thresholds of z>2.0. (**C**) Preferred theta-band phase of gamma-band amplitude for channel-frequency pairs in significant clusters in B. (**D**) Polar histograms of preferred CSD theta-band phase of gamma-band amplitude, separated by slow and fast gamma, pooled across recordings; black line represents median phase.

We used *P*_episode_ (20–22) to detect bouts of theta-band oscillations during task performance. Because our aim was to compare the absolute prevalence of theta-band oscillations across recording depths, we applied a common power threshold to all channels rather than a channel-specific threshold. Accordingly, rather than fitting the chi-square background model to each recording contact individually, we computed a grand-average power spectrum across all channels for each recording and fit the background model to this common spectrum. The 95th percentile threshold derived from this fit was then applied uniformly across channels. This approach allowed us to identify epochs where theta-band power exceeded a common absolute level, rather than a channel-specific relative level, enabling comparison of the absolute prevalence and duration of theta bouts across recording depths. Figure 2B depicts a representative theta bout, localized to a subset of adjacent recording sites. We also observed a systematic laminar organization of phase of the theta-band oscillation, with an approximately 180-degree reversal near the CA1 pyramidal cell layer (Fig. 2C). Theta bout prevalence (*P*_episode_) and bout duration closely mirrored the laminar profile of theta-band power (Fig. 2A, 2D).

Having characterized theta bouts across recording depths, we asked whether their laminar distribution aligned with the physiological and anatomical landmarks identified above. The theta reference channel was consistently located superficial to the CA1 pyramidal cell layer reference (Fig. 2A), at the depth corresponding to the *stratum lacunosum-moleculare* as independently identified from the ripple- and DS-evoked CSD reversals. Thus, the theta-band power maximum provided a third, independent physiological landmark that converged with the SWR- and DS-based references. Consistent with observations in rodents, the highest theta-band power localized to CA1 *stratum lacunosum-moleculare* (10, 11).

LFP power above 20 Hz also varied systematically with depth, most prominently in CA1, where power spectra consistently showed a peak in the gamma band (35-120 Hz; Fig. 3A). In rodent CA1, gamma-band amplitude has been shown to vary with the phase of the ongoing theta-band rhythm (23), so we next asked whether gamma-band amplitude at these depths was theta-modulated and whether the preferred phase of that modulation differed across gamma frequencies. We computed phase-amplitude coupling between the CSD-derived theta-band phase (3-7 Hz) on the theta reference channel and LFP amplitude from 35 to 120 Hz for every CA1 channel (see Methods). Throughout, theta-band phase is referenced to the 3-7 Hz CSD, with 0 corresponding to the theta peak and ±180° (±π) to the trough; positive phases therefore fell on the descending limb (peak to trough) and negative phases on the ascending limb (trough to peak). Phase-amplitude coupling was significant across a broad range of amplitude frequencies, but the preferred theta-band phase differed between gamma bands (Fig. 3). The amplitude of fast gamma (60-120 Hz) peaked near the theta-band trough, consistently across recordings (circular mean 162°, R = 0.94, n = 12 recordings, Rayleigh p = 8 × 10^-7^). Slow gamma (35-60 Hz) amplitude peaked later in the theta cycle, just past the trough on the ascending limb (circular mean −142°, R = 0.57, Rayleigh p = 0.016). The offset between bands averaged 56° and was positive (slow gamma later than fast gamma) in 10 of 12 recordings; the remaining two recordings showed the opposite ordering. This offset was significant across recordings (paired permutation test on preferred phase, p = 0.031; sign test on offset direction, p = 0.039).

The analyses so far characterized laminar structure through LFP signatures. We next turned to the single units recorded across these layers and asked how distinct cell types (Fig. 4A) are distributed along the depth axis. We pooled all units across the 12 sessions after aligning each session’s depth axis to the DS sink (0 µm; Fig. 4B-D; see Fig. S1 for individual-session distributions). The laminar distribution of units merged across sessions recapitulated hippocampal cytoarchitecture (Fig. 4B): principal cells were enriched in the DG granule cell layer and the CA3 and CA1 pyramidal cell layers, whereas interneurons were distributed more broadly across all layers. Burst propensity (defined as the ratio of short- to long-latency autocorrelogram firing (24)) and theta-band modulation (defined as the peak-trough asymmetry of autocorrelogram firing at theta frequencies (25)) varied systematically with depth as well, with the most strongly bursting and theta-modulated units concentrated in the same principal cell layers (Fig. 4C-D, Fig S1).

**Figure 4:**
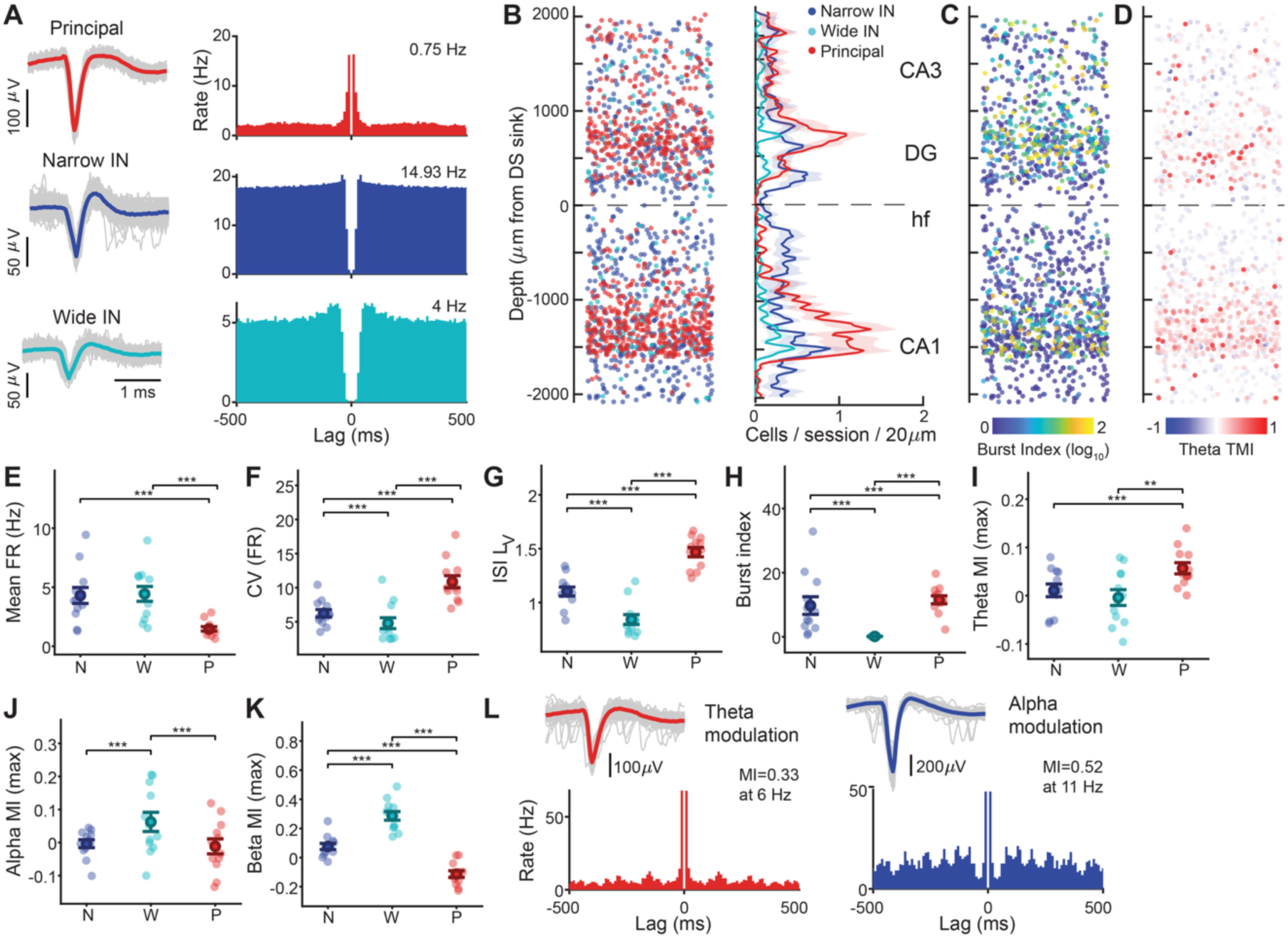
Laminar organization of hippocampal cell types, along with spiking and oscillatory properties. (**A**) Left: Three representative units, one per classified cell type: narrow-waveform interneuron (blue), wide-waveform interneuron (cyan), and principal (red). Right: Autocorrelogram (ACG) for each unit, 10 ms bins; ±500 ms lag. Mean firing rate is indicated at the top right of each ACG. (**B**) Left: putative cell type of every unit (1651 units, 12 sessions, 2 animals) plotted against depth, aligned to the dentate spike sink (0 µm); positive = above, negative = below, horizontally jittered for visualization; color as in (A). Right: unit counts per cell type and session in 20-µm depth bins (Gaussian-smoothed, 100 µm). Shaded area indicates SEM. Approximate subregions are shown on the right. hf: hippocampal fissure. (**C**) The same units as in (B), colored by burst index (log_10_-transformed). (**D**) The same units as in (B), colored by the maximum theta-band modulation index (theta TMI). (**E-H**) Distributions of single-unit spiking metrics by cell type. Small dots show each session’s mean; the large marker and error bar show the mean ± SEM across sessions. N, narrow-waveform interneuron; W, wide-waveform interneuron; P, principal cell. (**E**) Mean firing rate. (**F**) Coefficient of variation of the firing rate. (**G**) Local variation of the inter-spike interval (ISI LV). (**H**) Burst index (log_10_-transformed). (**I**) Maximum modulation index in the theta-frequency band (3-7 Hz) (**J**) Maximum modulation index in the alpha-frequency band (8-12 Hz), and (**K**) Maximum modulation index in the beta-frequency (13-30 Hz) band. (**L**) One sample unit exhibiting theta-band rhythmicity and one unit exhibiting alpha-band rhythmicity, colored by cell type as in (A), shown with the same waveform and ACG conventions as in (A). Maximum modulation index (MI) and its corresponding frequency are indicated at the top right of each ACG. Brackets in (E-K) denote pairwise contrasts from linear mixed-effects models (metric ∼ cell type + (1|session)); p-values are FDR-corrected across all pairwise comparisons and metrics (*p < 0.05; **p < 0.01; ***p < 0.001).

We characterized the intrinsic firing properties of the three cell types across the recorded population (n = 1651 units within ±2000 µm of the dentate spike sink, 93.0% of all isolated units; 967 from monkey C and 684 from monkey M; mean ± SD per session: 57.8±26.5 narrow-waveform interneurons, 14.3±6.4 wide-waveform interneurons, 65.6±17.6 principal cells). Cell properties were compared across cell types with a linear mixed-effects model including a random intercept for session, with significance assessed by likelihood-ratio test. Between-session variance was small throughout (ICC ≤ 0.09, see Methods), indicating that these differences reflect consistent cell-type properties rather than session-to-session variability.

Principal cells fired sparsely, irregularly, and in bursts: they showed the lowest mean firing rate, the highest coefficient of variation (CV) of firing rate and local variation (LV) in the inter-spike interval (ISI) variability, and the strongest burst firing of any cell type (Fig. 4E-H, Fig. S2, all comparisons involving principal cells p<0.001). Together with their localization to the principal cell layers, these firing properties supported their classification as putative hippocampal principal cells. Interneurons, by contrast, fired more frequently and more regularly, consistent with a role as sustained inhibitory pacemakers. The two interneuron subtypes were themselves distinguishable: wide-waveform interneurons showed the least bursting and the lowest CV and local ISI variability of the three cell types, suggesting they form a distinct, more tonically firing inhibitory subclass.

Oscillatory properties further separated the three cell classes (Fig. 4I-L, Fig. S2). Wide-waveform interneurons showed the strongest modulation of the three classes in both the alpha- and beta-frequency bands (both p<0.001, Fig. 4I-K, Fig S2). Theta-band modulation showed the opposite tendency: it was strongest in principal cells and weaker in both interneuron classes, which did not differ from each other (narrow vs. wide, p=0.240, n.s.; each vs. principal cells, p≤0.01; Fig. 4I). These results reveal a graded shift across cell classes: overall, principal cells tended to be strongly entrained to theta, whereas wide-waveform interneurons showed stronger modulation at alpha and beta frequencies.

Finally, we examined the fine-timescale functional interactions between simultaneously recorded neurons using spike-time cross-correlograms (CCGs, Fig. 5). Across the 12 recording sessions, 113,827 unit pairs had enough coincident spikes in their cross-correlogram and were considered for further analyses (see Methods). Of these, 1380 (1.21%) showed a significant short timescale interaction, indicating that the detection of fine timescale coupling was sparse within the recorded units (Fig. 5A-B). Of these coupled neuron pairs, 68.3% had near-zero lag synchrony consistent with a common input, whereas the remaining 31.7% had asymmetric CCG peaks, consistent with synaptic-like interactions (Fig. 5C) (26, 27).

**Figure 5:**
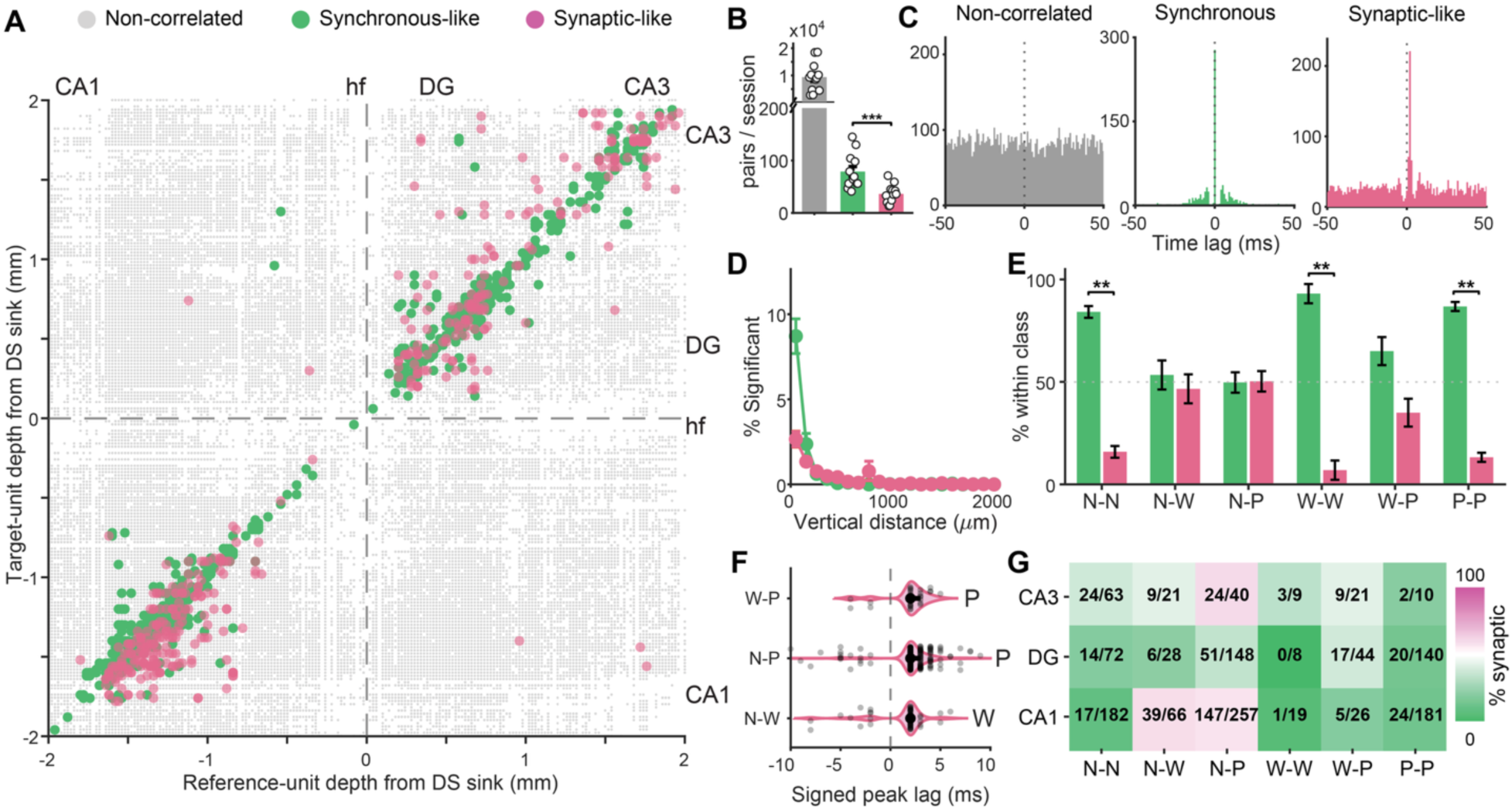
Fine-timescale spike-timing interactions reveal distance- and cell-type-dependent coupling. (**A**) Depth scatter of all unique unit pairs, pooled across sessions and aligned to the dentate spike (DS) sink of each recording. Each point represents one unique pair, plotted at reference-unit depth and target-unit depth relative to the alignment reference (dashed crosshair at 0 µm). Small gray points: non-significant valid pairs (see Methods and Results); Green: synchronous. Magenta: synaptic-like. Points falling near the diagonal indicate pairs with similar depths. (**B**) Mean number of unique pairs per session, by type of interaction, with individual session values overlaid (open circles), same color map as in (A). (**C**) Representative cross-correlograms (coincident spikes vs. time lag) for a non-correlated pair (left), a synchronous pair (middle), and a synaptic-like pair (right). (**D**) Significant interaction rate (% of valid unique pairs) as a function of vertical inter-unit distance (100 µm bins, up to 2000 µm), mean across sessions, for synchronous (green) and synaptic-like (magenta); errorbars represent SEM. (**E**) Cell-type dependence of interaction classification, mean across sessions. Asterisks: FDR-corrected significant difference between synchronous and synaptic-like interactions (**q < 0.01), and nonsignificant comparisons are left unannotated; errorbars represent SEM. (**F**) Signed peak-lag distributions of synaptic-like cross-type pairs. Each point represents one significant synaptic-like pair whose two units are of different cell types; positive lag indicates that the labeled cell type (right of each violin) led the interaction. Violins show the per-pair lag distribution (Gaussian kernel density). W-P, Principal leads; N-P: Principal leads, N-W: Wide-waveform interneuron leads). (**G**) Fraction of significant interactions that are synaptic-like vs. synchronous by subfield (rows) and cell-type comparison (columns; N = Narrow-waveform interneuron, W = Wide-waveform interneuron, P = Principal), for pairs with both units in the same subfield. Color encodes the fraction (green = 0%, magenta = 100%); text gives pooled counts across sessions, n(synaptic-like)/n(total).

The probability of detecting fine-timescale functional coupling decreased markedly with increasing spatial separation between units, with the odds declining by ∼65% per additional 100 µm for synchronous interactions and ∼30% for synaptic-like interactions (per-session binomial logistic regression, each slope<0 across sessions). This distance dependence was significantly steeper for synchronous coupling (Fig. 5D; paired Wilcoxon signed-rank test comparing session-level logistic regression slopes, p < 0.001).

We next asked whether the balance between synchronous and synaptic-like interactions depended on the cell-type identity of the two units involved (Fig. 5E). Pairs composed of two units of the same putative cell type were strongly biased toward synchronous rather than synaptic-like interactions: narrow-waveform interneuron pairs (N-N: 84.1 ± 2.9% synchronous; paired Wilcoxon signed-rank test across sessions, W = 78, p = 4.9×10⁻⁴, BH-FDR q = 1.5×10⁻³), wide-waveform interneuron pairs (W-W: 93.1 ± 4.7% synchronous; W = 45, p = 3.9×10⁻³, q = 7.8×10⁻³), and principal-principal pairs (P-P: 86.8 ± 2.2% synchronous; W = 78, p = 4.9×10⁻⁴, q = 1.5×10⁻³) each showed a significant excess of synchronous over synaptic-like classification. By contrast, none of the three cross-type combinations showed a significant departure from an even split between classes (narrow-waveform interneuron vs. wide-waveform interneuron, N-W: 53.4 ± 7.1% vs. 46.6 ± 7.1%, q = 0.91; narrow-waveform interneuron vs. principal, N-P: 49.7 ± 5.0% vs. 50.3 ± 5.0%, q = 0.91; wide-waveform interneuron-principal, W-P: 65.0 ± 6.8% vs. 35.0 ± 6.8%, q = 0.097).

The signed peak lags of synaptic-like pairs revealed a consistent directional hierarchy among cell types (Fig. 5F). For every cross-type combination, one cell type led the interaction (i.e., its spikes preceded the partner unit’s) far more often than expected by chance. Principal cells led narrow-waveform interneuron partners in 92.2% of synaptic-like N-P pairs (n = 232 pairs; Wilson 95% CI, 88.1–95.0%; one-proportion binomial test against chance, p < 10⁻¹⁰, BH-FDR q < 10⁻¹⁰) and led wide-waveform interneuron partners in 87.5% of W-P pairs (n = 32; 95% CI, 71.9–95.0%; p = 2.2×10⁻⁵, q = 2.2×10⁻⁵). Wide-waveform interneurons, in turn, led narrow-waveform interneuron partners in 81.0% of N-W pairs (n = 58; 95% CI, 69.1–89.1%; p = 2.3×10⁻⁶, q = 3.4×10⁻⁶). Together, these results are consistent with a directional hierarchy in the local circuit, with principal neurons driving local interneuron populations in CA1 (28) and extend this relationship to a specific hierarchy among interneuron subtypes.

To determine whether this cell-type organization was uniform across the hippocampal formation, we resolved the synchronous vs synaptic-like composition of significant pairs per hippocampal subfield. Because several cross-type combinations in some subregions were supported by few significant pairs, we report these results in Fig. 5G without statistical comparisons. The overall pattern was broadly preserved within each subfield: same-type pairs remained predominantly synchronous across CA1, DG and CA3, whereas N-P pairs were the most enriched for synaptic-like interactions, especially in CA1 and CA3. These results point to the value of integrating fine-timescale spike interactions with laminar markers of network activity when characterizing hippocampal circuit organization.

## Discussion

This study represents the first *in vivo* physiological characterization of the full laminar depth of the hippocampus in primates. Sharp-wave ripples and dentate spikes produced stereotyped current-source density profiles that provided physiological landmarks for CA1 and dentate gyrus, respectively, while theta-band activity showed a distinct depth dependence with maximal power and bout prevalence near CA1 *stratum lacunosum-moleculare*. When recordings were aligned across sessions using these physiological landmarks, putative principal cells and interneurons occupied distinct laminar distributions and differed systematically in their firing and oscillatory properties. Fine-timescale interactions were sparse and predominantly local but showed a reproducible cell-type structure, including a directional tendency in which principal cells led interneurons. Together, these findings establish a physiological framework for resolving hippocampal layers and local circuit organization in the awake primate brain.

Dense laminar coverage was important not only for anatomical registration but also for identifying the local generators of hippocampal field events. High-density recordings across CA1 layers have been proposed as the gold standard for distinguishing SWRs from other high-frequency activity, because their interpretation depends on the joint laminar organization of the sharp wave, ripple, and associated transmembrane currents (17). Our recordings therefore provide information that cannot be recovered from a high-yield recording restricted to the cell-body layer. Previous laminar recordings in macaque CA1 have shown that SWR-associated current reversals and cell-type distributions reveal a superficial-deep circuit organization (6). Our results extend this physiological characterization beyond CA1 by combining independent SWR and DS landmarks to resolve the CA3-DG-CA1 axis within the same penetration. Importantly, layer assignment did not depend on any single electrophysiological feature. Instead, multiple, independent signatures converged on the same laminar identities. SWR- and DS-triggered CSDs yielded complementary laminar signatures that converged with changes in single-unit density and the laminar profile of theta-band activity. The correspondence of these signatures with the well-characterized laminar physiology of rodent hippocampus (8, 10) suggests conservation of core circuit motifs despite substantial differences in hippocampal scale and cytoarchitecture (2). Together, these physiological landmarks provide a framework for registering recordings to the anatomy of the primate hippocampus when no single electrophysiological signal is sufficient to define anatomical boundaries.

Dentate spikes are also of interest beyond their utility as physiological landmarks. In rodents, DSs constitute a major hippocampal population event, distinct from SWRs, that can coordinate activity across the hippocampus and contribute to offline memory processing (29). The presence of a conserved DS laminar CSD in macaques therefore raises the possibility that DSs are part of a conserved network event across species and provides a physiological basis for testing their role in recognition memory in primates. A recent conference report has also described DSs in sleeping macaques (30), but a full characterization of DSs in primates is currently lacking. This gap may partly reflect the technical difficulty in identifying DSs in primates, for which dense laminar sampling has only become available recently.

The rate of both DSs and SWRs in primates differed from that reported in rodents. We found that SWRs occurred at around 13-15 events/min, which is within the broad range reported in awake primates (17, 31, 32). Rodent estimates are variable, but can be substantially higher during waking immobility and NREM sleep (7). We found that DSs occurred relatively infrequently during task performance (∼1.5 events/min). This rate lies within the broad range originally reported in rodents (0.6-30 events/min) (8) but it is substantially lower than the ∼15-30 events/min typically observed during rodent immobility and sleep/rest in recent studies (29, 33). Consistent with a contribution of behavioral state, preliminary macaque sleep recordings reported higher rates of both SWRs (∼19 events/min) and DSs (∼5.5 events/min) (30). Because DSs and SWRs depend strongly on behavioral state (7, 8, 29, 33), and reported SWR rates are additionally sensitive to detection criteria (17), differences in absolute rates cannot yet be interpreted as species differences.

Theta-band oscillatory activity was the third physiological signature that varied systematically with depth in our recordings. In primates, theta-band oscillations have been previously described as intermittent, in contrast to the sustained theta activity typically associated with rodent exploration (5, 12–14, 20, 34). Our results add a spatial dimension to this distinction: when theta bouts occurred, they were not distributed uniformly across the hippocampus but were preferentially expressed around CA1 *stratum lacunosum-moleculare* (10, 11). Theta-band oscillations also showed the characteristic phase shift across hippocampal layers described in rodents, further pointing to a conserved laminar organization. Thus, differences in the temporal expression of theta-band oscillations across species may coexist with conservation of its spatial and laminar organization.

Theta-band oscillations also organized higher frequency activity within CA1. Gamma-band amplitude was modulated by theta-band phase, with slow and fast gamma-band amplitudes peaking at different phases of the theta cycle. Fast gamma peaked near theta-band trough, while slow gamma was shifted to the ascending phase. This frequency-dependent phase separation resembles a canonical feature of rodent CA1, where slow and fast gamma are differentially coupled to theta and have been associated with different afferent pathways (23, 35, 36). The phase relationship of slow gamma in our recordings, however, differed from that typically reported in rodents, where slow gamma is strongest on the descending phase of the pyramidal-layer referenced theta. That said, direct comparison of absolute phase across studies is complicated by differences in theta reference and recording depth: as discussed above, theta undergoes a pronounced phase shift across CA1 laminae, and depth-resolved recordings have shown that preferred coupling phase can vary accordingly (37).

Our findings also differ from Abbaspoor et al., 2023, which reported little evidence for theta-gamma phase-amplitude coupling in macaque CA1, although theta was coupled to activity in the higher 95-150 Hz range (15). Their recordings, however, were confined to the CA1 pyramidal cell layer. Here, we found that theta-gamma coupling was strongest in the dendritic layers, where CA3 and entorhinal afferents terminate, suggesting that differences in laminar sampling may contribute to this discrepancy.

Together, these comparisons underscore that theta-gamma coupling is strongly shaped by recording location within CA1. Laminar differences in theta phase and gamma expression can influence both the apparent strength and preferred phase of coupling, complicating comparisons across studies. Additional variability may arise from behavioral state, frequency ranges used to define gamma bands (38) and species-specific circuit dynamics. Resolving how these factors shape theta-gamma coupling will require further investigation across species and behavioral states.

The laminar organization evident in the field potentials was accompanied by systematic differences among extracellularly defined cell classes. Putative principal cells were concentrated in the expected principal cell layers, and exhibited low firing rates, irregular firing, and strong burst propensity, while interneuron populations were distributed more broadly across layers. This combination of sparse firing, high bursting, and localization to principal cell layers recapitulates the canonical physiological phenotype of hippocampal pyramidal cells previously observed in macaque CA1 (6). Our results further revealed substantial functional heterogeneity within the inhibitory population. Narrow- and wide-waveform interneurons differed not only in the extracellular features used for classifying them, but also in their burst propensity, spike-train variability, and their modulation by network rhythms. These results support a distinction between the two defined interneuron groups, which is consistent with the extensive interneuron diversity found in rodents (2, 39–42) and with previous macaque recordings in which extracellularly defined inhibitory groups differed in firing statistics and oscillatory modulations (6). At the same time, while extracellular features in action potentials can capture reproducible functional diversity, they do not map uniquely onto transcriptomic or morphologically defined identities. Rather, narrow- and wide-waveform groups likely capture a broad subdivision within the much larger continuum of hippocampal inhibitory cell diversity (2).

Fine-timescale spike interactions provided a complementary view of this organization. Significant interactions were rare and decreased strongly with inter-unit distance, indicating that detectable fast coupling was predominantly local. The temporal structure of these interactions depended strongly on cell class: same-type pairs were biased toward synchronous interactions, consistent with shared input or other mechanisms of fast local synchronization, while cross-type pairs contained a larger fraction of asymmetric, synaptic-like interactions. Among cross-type synaptic-like pairs, principal cells tended to precede interneuron classes, matching the direction of principal-to-interneuron interactions in rodent CA1 (27, 28). Wide-waveform interneurons in turn tended to precede narrow-waveform interneurons, revealing a potential order within the inhibitory population. Although this asymmetry does not establish direct inhibitory connectivity, it provides additional evidence that the two defined interneuron classes belong to different positions within the local circuit. Previous macaque CA1 recordings have likewise shown that short timescale interactions between pyramidal and inhibitory populations depend on both pyramidal cell sublayer and inhibitory cell groups (6). Our results extend this organization across CA1, CA3, and DG, where the predominance of synchronous interactions and enrichment of the asymmetric principal- interneuron interactions were broadly preserved, suggesting a common organizational principle for local excitatory-inhibitory organization in the primate hippocampus.

Although the number of significant pairwise spike interactions was insufficient to resolve subfield-specific and cross-regional directionality, this sampling constraint highlights the complementary value of a laminar framework that combines spike and LFP signatures. Pairwise interactions require the simultaneous sampling of neurons that are functionally coupled, making such interactions relatively sparse even in large-scale extracellular recordings (27, 28). In this respect, laminar LFP and CSD signals provide a robust circuit-level readout.

In this study, recordings were obtained from two animals and sampled a restricted range of anterior-posterior and mediolateral positions in the hippocampus. Accordingly, extending this framework across the anterior-posterior and mediolateral axis, ideally with simultaneous recordings from multiple probes, will be important for understanding how laminar signatures vary systematically with recording position.

Beyond providing a physiological characterization across hippocampal layers and subregions, the framework established here opens the way to a new generation of experiments that can combine laminar recordings with pathway-specific perturbations in the primate. Because major hippocampal pathways terminate in distinct dendritic compartments, for example, CA3 Schaffer collateral input in CA1 *stratum radiatum* and entorhinal input onto more distal dendrites in the *stratum lacunosum-moleculare*, laminar recordings provide a direct means of resolving how pathway-specific perturbations reshape transmembrane currents and downstream neuronal activity. Combining high-density laminar recordings with selective manipulation of entorhinal, nucleus reuniens, or other afferent pathways could therefore link input-specific synaptic activity to oscillations and cell-type responses within the same local circuit, moving this framework from anatomical registration and cytoarchitectonic characterization towards a mechanistic account of primate hippocampal computation.

## Methods

All procedures were approved by the Institutional Animal Care and Use Committee of the Washington National Biomedical Research Center (protocol number: 4316-01) and adhered to the guidelines of the National Institutes of Health (NIH) for the care and use of laboratory animals.

### Subjects

Data were acquired in two female adult macaques (*Macaca mulatta*, monkey M average weight 6.8 kg, 8 years old; monkey C average weight 8.8 kg, 14 years old).

### Surgical procedures and electrophysiological recordings

Monkeys were implanted with a titanium headpost (Gray Matter Research) and a single recording chamber (2 cm diameter) positioned over the right hemisphere. Stereotaxic coordinates for chamber placement were determined from magnetic resonance imaging (MRI) scans obtained before implantation of the recording chamber, with the center of the chamber positioned at 9 mm anterior and 13 mm lateral to ear bar zero (EBZ) in monkey M, and 10 mm anterior and 13.5 mm lateral to EBZ in monkey C.

Five recording sessions were acquired for monkey C with the Value-Based Foraging Task, 3 sessions for monkey C with the Picture Viewing Task, and 4 sessions for monkey M with the Picture Viewing task. For all recordings, Neuropixels 1.0-NHP probes (IMEC, Leuven) were back loaded into metal guide tubes, with guide tube length set to reach approximately 10 mm below the dura. Neuropixels probes were slowly advanced using an oil hydraulic micromanipulator (Narishige Scientific Instruments). Extracellular electrical potentials from 384 channels were sampled at 30 kHz and streamed to a computer through the Neuropixels acquisition module (IMEC, Leuven) within a PXIe system (National Instruments, Austin, TX). Data acquisition was controlled by Open Ephys software (v0.6.7; www.open-ephys.org) and saved in binary format. Raw data were filtered and saved separately to an action potential band (AP band, 0.3-10 kHz, 30k Hz) and a local field potential band (LFP band, 0.5-500 Hz, 2.5 kHz). Behavioral analog signals were acquired simultaneously via a National Instruments DAQ device (PXIe-6341) at 40 kHz and synchronized with neural data through a shared TTL sync line. Recordings started around 45-60 minutes after probe insertion to allow for signal stabilization.

### Probe localization

On each recording day, a Neuropixels probe was initially advanced based on MRI planning until the hippocampus was reached. The presence of neural activity was monitored by the sound of the amplified signal of the deepest bank of contacts of the probe connected to an audio monitor and by observing spiking activity and local field potentials on a computer screen. We assessed the consistency of transitions between different brain regions such as the caudate, putamen and ventricles that lie above the hippocampus and compared with that expected from a brain atlas (43) as well as from preceding recording sessions.

### Behavioral presentation and task monitoring

A calibration task, a picture viewing task and foraging tasks were programmed in Unity3D and projected onto a large screen (160 x 90 cm, 1920x1080 pixels, 45 degrees vertical field of view) at 120-144 Hz. Avatar position was collected continuously at the same sampling rate. Bilateral eye position was monitored with an infrared eye-tracking system (EyeLink 1000, SR Research). Rewards were delivered via a pump-controlled spout.

### Picture Viewing Task

Monkeys were seated in front of a projector screen and presented with natural images subtending 24x32 degrees of visual angle. On each trial, an image remained available for free viewing until the monkey accumulated 7 seconds of viewing time. Each block consisted of 16 novel images, followed by a second presentation of the same images. Behavioral sessions comprised 6-9 blocks, corresponding to 96-144 unique images per session, with a new image set used in each session. No reward was delivered during image-viewing trials. Monkeys instead received food-slurry rewards for successful performance on interleaved, non-image eye-calibration trials. These trials began with the appearance of a small white fixation spot at pseudorandomly selected locations on the display. Monkeys were required to fixate on the spot and hold their gaze within a 2-degree window after which the color of the fixation spot switched from white to yellow and reward was delivered.

### Value-Based Foraging Task

Monkeys were trained to use a joystick (Logitech Extreme 3D Pro) to navigate in a virtual arena, from the perspective of a first-person avatar. At the start of each trial, the avatar was placed at a random border location facing toward the arena center. Each trial included a set of 9 foraging targets scattered across the environment. Targets were associated with low, medium, or high reward values. Whenever monkeys harvested available targets, they received drops of food-slurry reward and audible reinforcement beeps that were linearly related to the amount of reward. The specific reward contingencies varied by task condition: in the color-value mapping, reward value was determined by the color of the target; in the spatial-value mapping, the color was the same for all targets and reward value was determined by the zone in which the target appeared. Each recording session comprised alternating blocks (10-20 trials each) of two distinct color-value mappings and one spatial-value mapping. Task order was randomized across sessions. On average, monkeys completed approximately 40 trials per mapping per session.

### Single unit analyses

#### Spike sorting

All spike sorting, curation, and quality control steps were performed using custom MATLAB and Python scripts in conjunction with standard Kilosort4 (44) and Phy2 workflows (https://github.com/cortex-lab/phy).

Spike sorting outputs were manually curated using Phy to merge or split clusters and remove noise or multi-unit activity based on waveform features, firing rates, and auto-/cross-correlograms. Following curation, clusters were subjected to quality metric (QM) thresholds, including sliding refractory period violations, noise cut-off and amplitude thresholds, to define a final set of high-confidence single units (45).

Across 12 sessions (Monkey C: n = 8; Monkey M: n = 4), Kilosort4 identified 5596 units (Monkey C: 4309; Monkey M: 1287). After manual curation in Phy and application of quality-control criteria, 1776 units (31.7% of all Kilosort-identified units) were retained as final high-quality single units: 1047 from Monkey C (mean = 130.9 ± 39.4 units/session; range: 82-195) and 729 from Monkey M (mean = 182.3 ± 26.0 units/session; range: 155-206).

### Waveform extraction

Mean spike waveforms were extracted from the filtered raw data for each good unit (200 waveforms per unit). For each unit, the channel with maximum waveform amplitude was identified, and the mean waveform on that channel was used for all subsequent waveform-based analyses. Waveforms were mean-subtracted, then aligned in time such that the dominant trough (for negative-going waveforms) or peak (for positive-going waveforms) fell at 30% of the extraction window, following the WaveMAP alignment protocol (46). Aligned waveforms were normalized by their maximum absolute amplitude prior to computing waveform-shape metrics.

### Waveform metrics and cell-type classification

Units were classified into three putative cell types: narrow-waveform interneurons, wide-waveform interneurons and principal cells based on waveform morphology and firing dynamics. Classification relied on two complementary features: trough-to-peak latency extracted from the average extracellular waveform and the rise time of the spike autocorrelation (ACG) histogram (47). In brief, autocorrelation histograms were computed for each unit and fitted with a triple exponential equation to quantify firing dynamics, supplementing classical waveform-based classification approaches. This combined approach leverages both the biophysical properties reflected in spike shape and the temporal firing patterns characteristic of distinct cell classes.

Narrow-waveform interneurons were identified by a trough-to-peak latency less than 0.425 ms. Wide-waveform interneurons were classified as units with trough-to-peak latency greater than 0.425 ms combined with an autocorrelation histogram rise time exceeding 6 ms. All remaining units were classified as putative principal cells.

### Single unit metrics

Spike times for each unit were binned into 100 ms bins to compute mean and coefficient of variation (CV) of the firing rate across the recording. Interspike interval (ISI) statistics were computed from the full spike train to measure local variation (LV) (48).

We quantified burst propensity as the ratio of the mean ACG rate at very short lags (3-5 ms) to the mean ACG rate at long lags (200-300 ms), providing a conservative baseline normalization that is less sensitive to overall firing rate (24).

To quantify rhythmic modulation of spiking, we computed frequency-resolved modulation indices from each unit’s spike-train ACG. Oscillatory modulation strength was quantified over a logarithmically spaced frequency grid (1/8-octave spacing) and summarized within three canonical frequency bands: theta (3-7 Hz), alpha (8-12 Hz), and beta (13-30 Hz). For each frequency within the band, we compared the mean ACG rate in a trough window to that in a subsequent peak window positioned according to the expected oscillatory period and defined a modulation index (25). For each band, we report the maximum modulation index within that band (a measure of how strongly spiking is rhythmically modulated at any tested frequency in that range) and the frequency at which that maximum occurred ("best frequency"). Because best-frequency estimates are only meaningful when a unit exhibits genuine rhythmic structure, best-frequency values were only considered for a given band, in both statistical analyses and figures, for units whose maximum modulation index in that band exceeded 0.1; units below this threshold were excluded from best-frequency comparisons (but not from comparisons of the modulation-index magnitude itself, which were computed for all units regardless of modulation strength).

### Statistical analysis for cell-types

All statistical comparisons across the three cell-type groups (narrow-waveform interneuron, wide-waveform interneuron, principal cell) were performed on the depth-filtered unit population described in the section ‘Anatomical depth alignment of single units’.

For each metric (firing rate, CV, LV, burst index, theta/alpha/beta max MI and best frequency), we fit a model of the form *metric ∼ cell_type + (1 | session)*, with recording session included as a random intercept to account for the non-independence of units recorded in the same session. Firing rate, firing rate CV, and burst index were log_10_-transformed prior to model fitting to satisfy approximate normality of model residuals, given their right-skewed distributions. Estimated marginal means and the intraclass correlation coefficient (ICC) were obtained from this model fit by restricted maximum likelihood (REML). ICC was calculated as the proportion of total variance attributable to between-session differences (session random-intercept variance divided by the sum of session and residual variance), to assess whether cell-type differences reflected consistent cell-type properties or were instead driven by session-to-session variability. Pairwise comparisons between cell types (narrow-waveform interneuron vs. wide-waveform interneuron, narrow interneuron vs. principal cell, wide interneuron vs. principal cell) were each assessed by likelihood-ratio test: for a given pair, we fit a reduced model in which the two cell types under comparison were merged into a single factor level while the third cell type remained distinct, and we compared this reduced model to the full three-level model via a likelihood-ratio test. We did not additionally test whether cell type had any effect overall, because the pairwise contrasts are the comparisons of interest.

The pairwise contrast p-values from every metric and every cell-type pair were pooled together and corrected once with Benjamini-Hochberg false discovery rate (FDR) correction, rather than corrected separately within each metric. Significance brackets on error bar and violin plots reflect these FDR-corrected pairwise contrasts (* p < 0.05, ** p < 0.01, *** p < 0.001).

### Anatomical depth alignment of single units

To analyze units across recording sessions and animals despite differing probe insertion depths, each unit’s anatomical position was expressed relative to a session-specific anatomical landmark (see below: Hippocampal layer identification) rather than as an absolute probe depth. Each unit’s depth was re-expressed as its distance from this landmark (depth aligned = unit depth − session landmark depth), and only units within ±2000 µm of the landmark were retained for statistical analysis and reported figures; this depth window was applied uniformly to every metric. Of the 1776 single units identified after spike-sorting curation, 1651 fell within this boundary (967 in monkey C, 684 in monkey M). The neurons that fell outside this boundary were recorded during deeper probe penetrations and likely include neurons in the deep layers of the underlying perirhinal cortex.

### Single-unit coupling

Fine-timescale coupling between simultaneously recorded units was quantified using spike-time cross-correlograms (CCGs) computed for all ordered pairs of units using 1 ms bins. To isolate millisecond-scale interactions from slower co-modulation in firing rate, CCGs were corrected using interval jitter. For each of 500 surrogate datasets, each spike was randomly reassigned to a time drawn uniformly within the same 25 ms interval as the original spike. This procedure preserved the number of spikes from each unit within each fixed 25-ms interval, while randomizing spike timing within the interval and thereby disrupting finer-timescale relationships. The expected CCG under this null model was obtained by averaging across the 500 jittered CCGs, and the jitter-corrected CCG was calculated as the observed CCG minus this average surrogate CCG.

Short-latency coupling was quantified within a ±10 ms window centered on zero lag. The zero-lag bins were retained in the analysis. The background level of each jitter-corrected CCG was estimated as the mean across two flanking windows (−100 to −50 ms and +50 to +100 ms). For each ordered unit pair, the coupling peak was defined as the maximum value of the jitter-corrected CCG within the ±10 ms window, and coupling strength was quantified as the difference between this peak and the pair-specific background level. Peak lag was defined as the lag corresponding to this maximum. Only pairs with more than 100 total coincidences in the raw CCG were considered valid and included in further analysis.

To determine whether a pair exhibited a significant fine-timescale interaction, the CCG was normalized by the geometric mean firing rate of the two units (√(FR_i × FR_j)). The standard deviation of the normalized CCG was calculated across the two flanking baseline windows. A peak z-score was then calculated as the baseline-subtracted, firing-rate-normalized peak amplitude divided by this standard deviation. Pairs with a peak z-score ≥ 7 were classified as significant interactions; all other pairs were classified as non-correlated.

The CCG asymmetry index (CA) was calculated from the baseline-subtracted, jitter-corrected CCG as the difference between summed counts at ±5 ms lags divided by their sum. Significant interactions were classified into two functional classes based on CCG asymmetry, peak lag, and peak width. Interactions were classified as synchronous if |CA| ≤ 0.3, |peak lag| ≤ 1 ms, or peak width was > 4 ms. All remaining interactions were classified as synaptic-like. These classifications describe CCG features consistent with putative common-input or directional synaptic interactions and do not imply anatomically confirmed connectivity.

Unit depth was taken from the position of the recording site on which the unit’s waveform was largest, converted to distance along the probe (20 µm site spacing, two sites per depth). Those values were aligned to the dentate spike sink before pooling sessions. Units were assigned to hippocampal subfields by this aligned depth: CA1, <0 µm; DG, 0 to 1200 µm; and CA3 1200 to 2000 µm. A pair was assigned to a subfield only when both units fell within the same subfield. Cross-subfield pairs, and pairs with either unit unassigned were excluded from within-subfield comparisons. Throughout, pair distance refers to the vertical separation of the two units along the probe.

Class proportions were computed within each session among significant interactions and compared across sessions with a paired Wilcoxon signed-rank test (Fig. 5B). The same session-level proportions were computed separately for each of the six cell-type combinations and compared the same way, with Benjamini-Hochberg correction across the six comparisons (Fig. 5E).

For each cross-type combination, the pooled fraction of synaptic pairs in which one of the cell types led was compared against chance (0.5) with a binomial test, with 95% confidence intervals (Fig. 5F). Because per-session counts within a single cross-type combination were small, this test pooled pairs across sessions.

#### LFP analysis

Unless otherwise specified, LFP analyses were performed in MATLAB using custom scripts and the FieldTrip toolbox (49) on one of the two interleaved columns of contacts. Subsampling to every 4th or 8th channel is noted where applied.

LFP signals were corrected for phase distortions introduced by the Neuropixels analog hardware bandpass filter by applying a time-reversal filtering procedure. The hardware filter was modeled as a first-order Butterworth bandpass filter with corner frequencies of 0.5 and 500 Hz. Signals were reversed in time, filtered using this model, and then reversed back, yielding phase-corrected LFP signals. Bad channels were defined as those with variance less than the median minus 3 SD and were replaced by interpolation from neighboring electrodes.

Segment epochs were defined relative to stimulus (Picture Viewing Task) or trial (Value-Based Foraging Task) onset, starting 500 ms after stimulus presentation to exclude any visually-evoked response.

Artifacts were detected from the mean LFP signal across channels and removed by excluding periods in which the signal exceeded the predefined global amplitude range, indicating railing artifacts, or showed large abrupt changes after 0.1-5 Hz bandpass filtering, indicating jump artifacts. Railing artifacts were removed with a 0.5 s pre-artifact and 1 s post-artifact buffer, whereas jump artifacts were removed with a 0.5 s buffer on both sides. Only artifact-free segments of at least 5 s were retained for analysis. Each segment was demeaned and linearly detrended to remove DC offsets and slow voltage drifts prior to spectral analysis.

Power spectra were estimated using the multitaper method (50) (DPSS tapers, 1 Hz smoothing) on 5-sec non-overlapping segments from 2 to 150 Hz. The aperiodic (1/f) component was removed using the FOOOF algorithm (19) with a fixed (no knee) aperiodic mode, fit on the median power spectrum across channels and subtracted in log space from each individual channel, yielding a corrected spectrum reflecting periodic power above the aperiodic background.

Bouts of theta activity in LFP data were quantified using an oscillatory episode detection algorithm that estimates the background power spectrum of the LFP in order to determine power and duration criteria (20). Artifact-free LFP segments were wavelet transformed using Morlet wavelets (51) (7 cycles) at 35 logarithmically-spaced frequencies between 2 and 38 Hz. For each recording, the background spectrum was estimated by fitting a linear regression in log-log coordinates to the grand-average power spectrum across all channels, assuming colored noise of the form *Af^-α^*. Because wavelet power values are distributed as *χ*^2^(2)(52), the fitted background power at each frequency represents the mean of this distribution. The power threshold, *P_T_*, was set to the 95^th^ percentile of the *χ*^2^(2) distribution at each frequency, and the duration threshold, *D_T_*, was set to 3 cycles. For each channel and frequency, all time intervals during which LFP power exceeded *P_T_* for a duration of *D_T_* were designated oscillatory episodes. Theta bouts were defined as contiguous time points containing oscillatory episodes at any frequency within the theta band (3-7 Hz). *P*_episode_ was defined as the proportion of time across all data segments during which both thresholds were exceeded. The use of a common threshold derived from the grand-average spectrum, rather than channel-specific thresholds, enabled comparison of absolute theta bout prevalence and duration across recording depths.

To construct the underlay in Figure 2C, the channel with the highest theta *P*_episode_ value was selected as the reference. Theta bouts on this channel lasting at least 1000 ms were each aligned to the peak of their second theta cycle, defined as the maximum of the band-pass–filtered (3-7 Hz) signal within the second cycle following bout onset. For each such bout, instantaneous theta phase was then extracted from every channel – including the reference – within a ±500 ms window around this common alignment point, as the angle of the analytic signal (Hilbert transform of the 3-7 Hz filtered LFP). Thus, all channels were sampled at the same set of reference-defined bout times, rather than at bouts detected independently on each channel. For each channel, phases were averaged across these bouts at each time lag by circular mean, computed as the argument of the mean unit phasor, 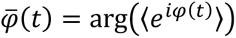. The resulting mean phase is displayed as a function of channel depth and time relative to the alignment point.

### Current source density

The current source density (CSD) was computed using the inverse CSD method with cubic splines, as implemented in the iCSD toolbox for MATLAB (53) on the LFP time series from every 4^th^ channel of one of the columns (80 µm spacing). A tissue conductivity of 0.3 S/m and an assumed current source diameter of 0.5 mm were used as model parameters. The resulting CSD was spatially smoothed with a Gaussian kernel (σ = 0.05 mm) and converted to units of µA/mm³. CSD was computed segment by segment and subsequently averaged across segments to produce the final laminar CSD profile.

### Dentate spikes

Dentate spikes (DS) were detected using the methods and scripts described in (54) and implemented in custom Python software. Briefly, we detected peaks of the filtered LFP (5-100 Hz, 4^th^ order Butterworth) in the channels spanning the dentate gyrus (DG). The detection channel was selected per session in the following way: candidate channels were first scanned and ranked by DS-triggered amplitude and event count, and the final channel for detection was confirmed by inspecting the DS-triggered CSD profile sink/source consistency. A threshold of 7 times the median absolute amplitude of the filtered LFP was employed, and we only considered events that were separated by at least 50 ms.

### Sharp-wave ripples

To identify the optimal reference channel for Sharp-wave ripple (SWR) events, we computed a ripple band score for each channel. This score was calculated by dividing the power in the ripple band (120-200 Hz) by the power in a broadband frequency range (70-300 Hz), using a Welch spectrum (4-s Hann windows overlapping by 50%). The channel with highest score was defined as the reference channel.

SWRs were detected on the selected channel using a previously described method (55), implemented in custom Python software. Briefly, the raw LFP of the reference channel (subsampled to 1,250 Hz) was filtered in the 120-200 Hz band (4^th^ order Butterworth) and transformed into a normalized squared signal (NSS). Peaks were identified by thresholding the NSS, merging neighboring events, and discarding events with excessive duration. Thresholds were computed as multiples of the standard deviation of the NSS: the peak power threshold was set to 5 standard deviations and the start/end cutoff to 2 standard deviations above the mean NSS. Duration limits were set to 15 and 100 ms, and events separated by fewer than 30 ms were merged prior to peak thresholding. The optimal detection channel within the CA1 pyramidal layer was confirmed by scanning all channels and comparing the one with the highest mean peak z-score across detected events with the channel selected above.

For each SWR event, time-frequency power (30-300 Hz, continuous Morlet wavelet transform (51)) was z-scored per-event against that event’s own pre-event baseline period, then averaged across events within a session. Each event’s peak ripple frequency (argmax of z-scored power in a ±20 ms window around its own aligned center) was extracted as a byproduct of this same computation and its distribution reported across all events.

### Phase-amplitude coupling

Phase-amplitude coupling (PAC) was computed between the phase of theta oscillations (3-7 Hz) extracted from the CSD and the amplitude of higher-frequency oscillations (35-120 Hz in 5 Hz steps) from the LFP. The CSD was computed on every 4^th^ channel of one of the columns (80 µm spacing). The 3 adjacent channels with the highest theta-band power were selected as the phase reference. Theta phase was extracted by bandpass filtering the averaged CSD signal across these 3 channels (3rd order Butterworth, zero phase) and applying the Hilbert transform to obtain the instantaneous phase at each sample.

Analysis of PAC was restricted to periods of high theta amplitude, defined as samples in which CSD theta amplitude exceeding the 50^th^ percentile of its distribution continuously for at least 0.25 s. For every 2^nd^ LFP channel of one of the columns (40 µm spacing) and each amplitude frequency (35-120 Hz; steps of 5 Hz; 10 Hz bandwidth), the signal was bandpass filtered, and the amplitude envelope was extracted via Hilbert transform. The modulation index (MI) was computed following (56) as the Kullback-Leibler divergence of the phase-binned amplitude distribution (18 bins of 20°) from a uniform distribution. Statistical significance was assessed by z-scoring the observed MI against a surrogate distribution of 500 MI values, each generated by circularly shifting the amplitude envelopes relative to the theta phase time series by a randomly chosen offset. Within each surrogate, the same shift was applied to all channel-frequency pairs, preserving the covariance structure across channels and frequencies. Significance was further assessed with cluster-based permutation testing (57) to control the family-wise error rate. Channel-frequency pairs exceeding a cluster-forming threshold of z > 2 were grouped into clusters, with adjacency defined by sharing an edge in the channel x frequency grid, and each cluster was summarized by the sum of its z-scores. A null distribution was built from 500 permutations; on each permutation the full surrogate procedure was repeated, clusters were formed by the same criterion, and the largest cluster statistic was retained. Observed clusters were considered significant at p < 0.05 if their cluster statistic exceeded the 95^th^ percentile of this maximum-statistic null distribution.

The preferred theta phase for each channel-frequency pair surviving cluster-based permutation testing was taken as the center of the phase bin with the highest mean amplitude in the 18-bin phase-amplitude histogram. Within each recording, analysis was restricted to channels at or below a boundary channel placed at the location closest to the putative CA1 *stratum radiatum*, and within 1200 µm ventral to that boundary. Clusters of significant coupling were defined separately within each band, with connectivity evaluated independently above and below 60 Hz so that clusters spanning the slow/fast gamma boundary were divided. Each contributing channel supplied a single preferred phase, taken at the amplitude frequency with the highest coupling z-score among frequencies significant for that channel.

Because channels within a recording share a phase reference and are spatially correlated, each recording was treated as a unit of analysis for all inferential statistics. For each recording and band, a circular mean was computed across contributing channels; group statistics were then computed across these recording-level means (n = 12). Distributions shown in Fig. 3D include all contributing channels, but all reported statistics are computed across recordings.

Mean direction and mean resultant length R were computed as the argument and modulus of the mean unit vector. Departure from uniformity was assessed with the Rayleigh test. The offset between bands was computed within each recording as the angular difference between its slow- and fast-gamma circular means, wrapped to (-180°, 180°]. Whether these offsets were rotated away from zero was tested by sign-flip permutation. Under the null hypothesis that offsets are symmetrically distributed about zero, the sign of each recording’s offset is exchangeable; the test statistic, the resultant component perpendicular to zero (|∑ sin(*d_i_*)|), was therefore recomputed for all 2^12^ sign assignments, and the p-value taken as the proportion of assignments equaling or exceeding the observed value. The consistency of offset direction was additionally assessed with an exact two-tailed sign test.

#### Hippocampal layer identification

Recordings were aligned across sessions and monkeys using a single anatomical anchor per session: the channel with peak current sink amplitude during the dentate spikes. This DS-sink channel served as the session-specific reference point for all depth alignment.

To confirm that DS-sink-anchored alignment corresponded to a consistent laminar position across sessions, we validated it against three independent physiological landmarks. First, we identified the channel with peak sharp-wave ripple power in the same CSD, which marks the CA1 pyramidal cell layer (58, 59). Second, from the single-unit population, we confirmed the CA1 pyramidal cell layer as the depth band containing the highest density of putative pyramidal cells (consistently the thickest such band across recordings) and confirmed that bursting neurons were likewise concentrated there, consistent with the expected laminar organization of CA1 (4, 6, 16, 58). Third, we computed the CSD from every fourth channel within a probe column (inter-site spacing 80 µm) and identified, per session, the three adjacent channels with the highest theta-band power; in rodents, such theta-associated current sinks provide a reliable landmark for *stratum lacunosum-moleculare* (10, 11). In every session, the theta-defined landmark, the ripple-power reference channel, and the unit-density-defined CA1 pyramidal layer occupied consistent positions relative to the DS-sink anchor and to one another. Together, these independent criteria confirmed that DS-sink alignment reliably provided a consistent laminar reference point across sessions and monkeys.

## Acknowledgements

We thank Megan Jutras for project support, Mary A. Rosu and Sada R. Nichols-Worley for technical assistance, and Autumn J. Mallory and Ryan Ressmeyer for guidance in the design of the Neuropixels drive adaptors. This work was supported by the Schmidt Science Fellows, in partnership with the Rhodes Trust (S.M.L.); Simons Foundation SFI-AN-NC- GB-Independence Postdoctoral-00006781 (S.M.L.), SCGB 542955 (E.A.B.) and SCGB NC-GB- CULM-00002730 (E.A.B); and by the NIH: NINDS-U19NS107609 (E.A.B.), NINDS- UF1NS126485 (E.A.B.), ORIP-P51OD010425 (WaNBRC), and an NIA training grant T32AG066574 (E.C.S.B.).

---

**Figure S1.**
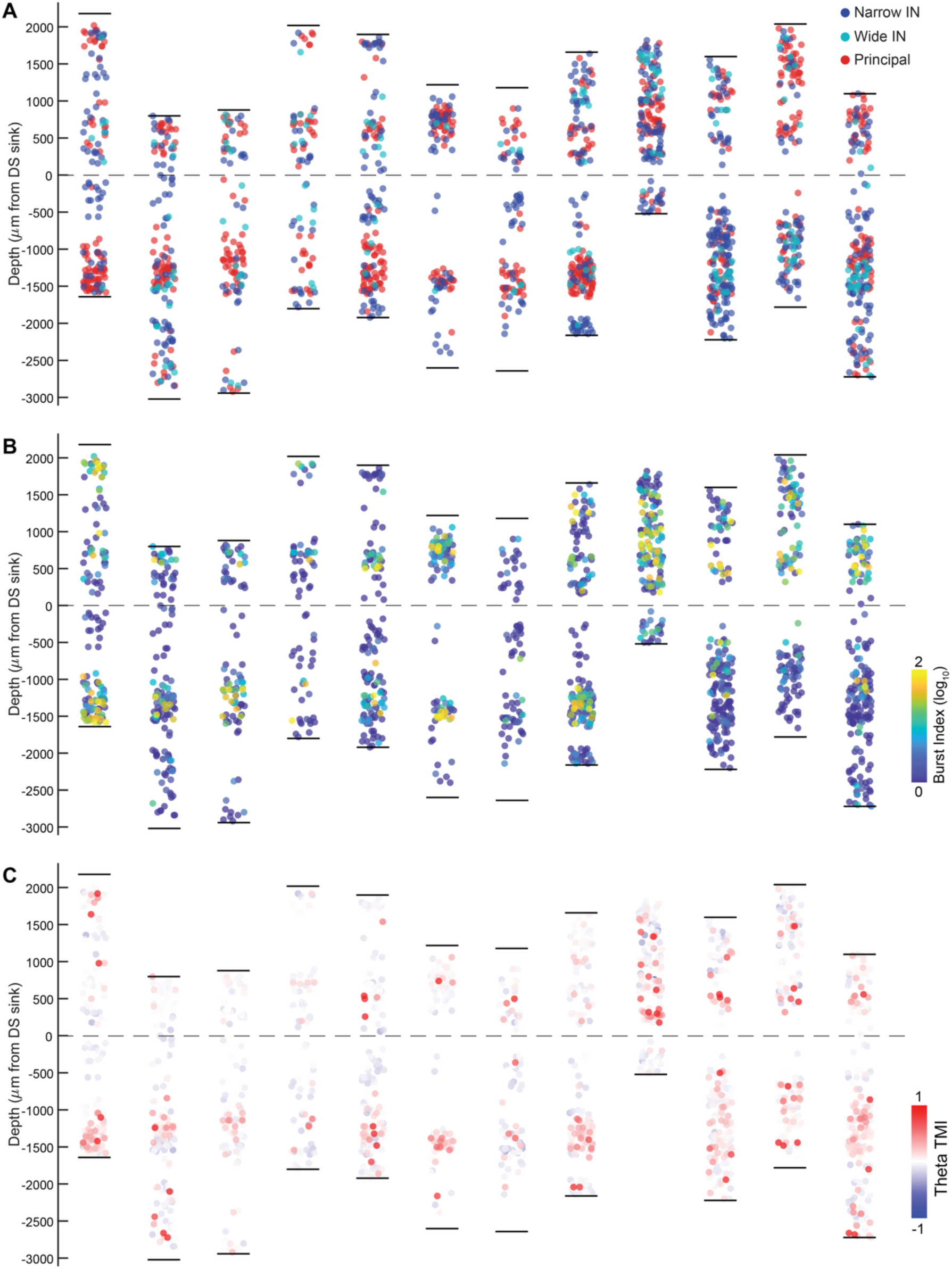
Per-session laminar distribution of cell types and spiking properties. Single-unit data shown separately for each recording session (N = 12 sessions from 2 monkeys). Sessions are ordered by animal and then by recording date. For all panels, each unit is plotted at its depth aligned to the dentate spike (DS) sink (0 µm; dashed line), with units jittered horizontally within each session’s column for visibility. Short horizontal ticks at the top and bottom of each column mark the dorsal and ventral extent of the recorded units along the probe for that session. The depth axis spans the full recorded range across sessions. (**A**) Putative cell type of each unit: narrow-waveform interneurons (Narrow IN, blue), wide-waveform interneurons (Wide IN, cyan), and principal cells (Principal, red). (**B**) The same units colored by their burst index (log_10_-transformed); color scale at the bottom right corner of the panel. (**C**) The same units colored by their theta modulation index (TMI); color scale at the bottom right corner of the panel. The diverging scale is centered at zero (no modulation), with positive and negative values in warm and cool colors, respectively.

**Figure S2.**
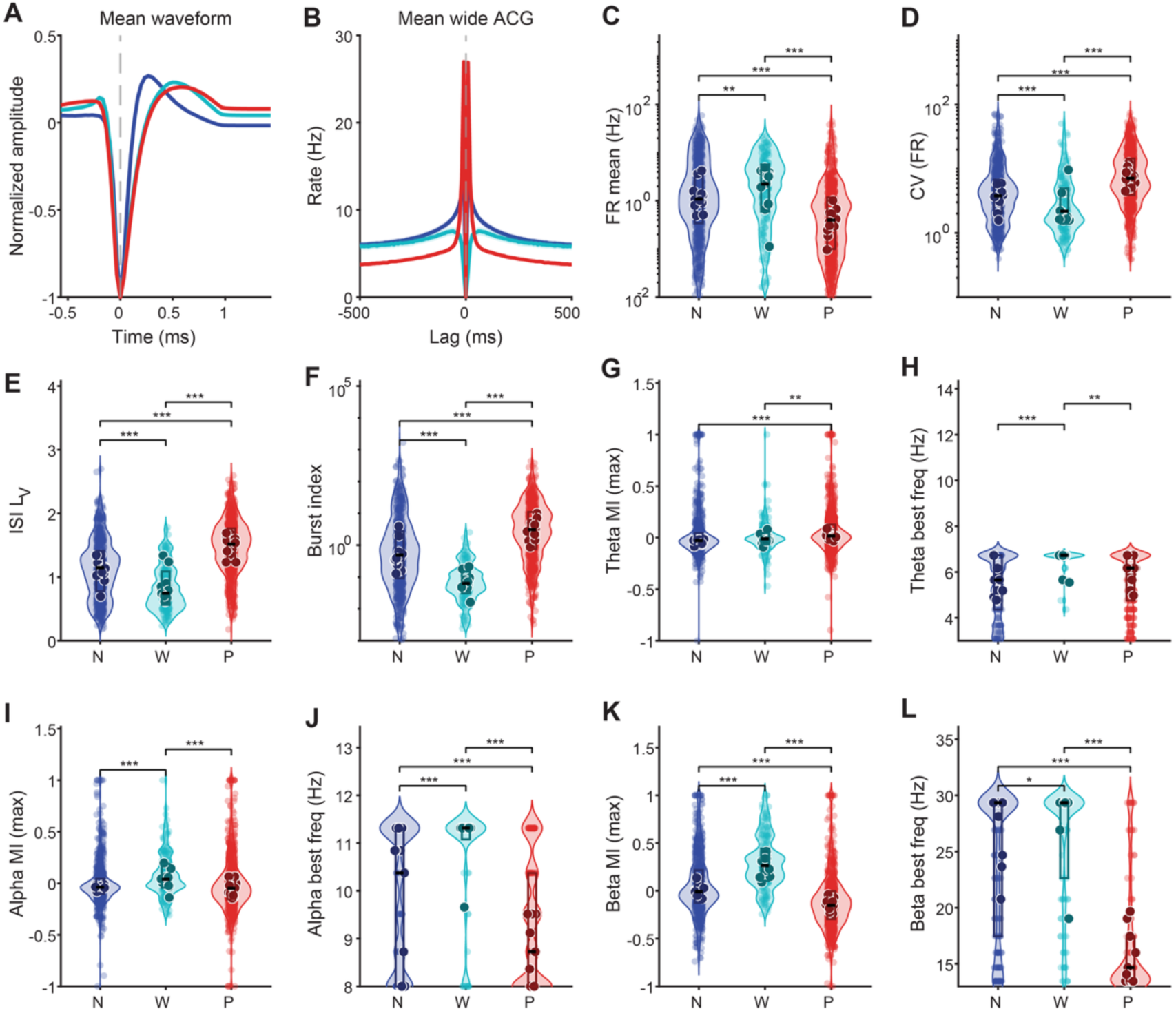
Spiking properties by cell type. Single units pooled across sessions and grouped into three putative cell types: narrow-waveform interneurons (Narrow IN, blue), wide-waveform interneurons (Wide IN, cyan), and principal cells (Principal, red). (**A**) Session-balanced mean spike waveform for each cell type, normalized to the trough; lines show the mean and shaded bands the SEM across session means. (**B**) Session-balanced mean autocorrelogram (ACG, 10-ms bins) for each cell type (mean ± SEM across session means); y-axis in spikes/s. (**C-L**) Distributions of single-unit spiking metrics by cell type, shown as violin plots (kernel density) with individual units overlaid as colored points, session means as darker larger points; and the central line marking the median and the box the interquartile range. (**C**) Mean firing rate. (**D**) Coefficient of variation of the firing rate. (**E**) Local variation of the inter-spike interval (ISI LV). (**F**) (log_10_ transformed). (**G**) Theta modulation index (maximum across 3-7 Hz). (**H**) Best theta frequency, restricted to units with theta modulation index > 0.1 (G). (**I**) Alpha modulation index (maximum across 8- 12 Hz). (**J**) Best alpha frequency, restricted to units with alpha modulation index > 0.1 (I). (**K**) Beta modulation index (maximum across 13-30 Hz). (**L**) Best beta frequency, restricted to units with beta modulation index > 0.1 (K). Colors are consistent across all panels. Brackets in (C-L) denote pairwise contrasts from linear mixed-effects models (metric ∼ cell type + (1|session)); p-values are FDR-corrected across all pairwise comparisons and metrics, pooled into a single family (*p < 0.05; **p < 0.01; ***p < 0.001).

## Notes

### Competing Interest Statement

The authors have declared no competing interest.

## References

1. D. G. Amaral, M. P. Witter, The three-dimensional organization of the hippocampal formation: a review of anatomical data. Neuroscience 31, 571–591 (1989).

2. D. G. Amaral, P. Lavenex, R. Insausti, “Hippocampal Neuroanatomy” in The Hippocampus Book, R. Morris, D. G. Amaral, T. Bliss, K. Duff, J. O’Keefe, Eds. (Oxford University Press, 2007), pp. 37–114.

3. H. Kondo, P. Lavenex, D. G. Amaral, Intrinsic connections of the macaque monkey hippocampal formation: I. Dentate gyrus. J of Comparative Neurology 511, 497–520 (2008).

4. H. Kondo, P. Lavenex, D. G. Amaral, Intrinsic connections of the macaque monkey hippocampal formation: II. CA3 connections. J of Comparative Neurology 515, 349–377 (2009).

5. N. J. Killian, M. J. Jutras, E. A. Buffalo, A map of visual space in the primate entorhinal cortex. Nature 491, 761–764 (2012).

6. S. Abbaspoor, K. L. Hoffman, Circuit dynamics of superficial and deep CA1 pyramidal cells and inhibitory cells in freely moving macaques. Cell Reports 43, 114519 (2024).

7. G. Buzsáki, Hippocampal sharp wave-ripple: A cognitive biomarker for episodic memory and planning. Hippocampus 25, 1073–1188 (2015).

8. A. Bragin, G. Jando, Z. Nadasdy, M. van Landeghem, G. Buzsáki, Dentate EEG spikes and associated interneuronal population bursts in the hippocampal hilar region of the rat. Journal of Neurophysiology 73, 1691–1705 (1995).

9. D. Dvorak, A. Chung, E. H. Park, A. A. Fenton, Dentate spikes and external control of hippocampal function. Cell Reports 36, 109497 (2021).

10. G. Buzsáki, J. Czopf, I. Kondákor, L. Kellényi, Laminar distribution of hippocampal rhythmic slow activity (RSA) in the behaving rat: Current-source density analysis, effects of urethane and atropine. Brain Research 365, 125–137 (1986).

11. J. Brankačk, M. Stewart, S. E. Fox, Current source density analysis of the hippocampal theta rhythm: associated sustained potentials and candidate synaptic generators. Brain Research 615, 310–327 (1993).

12. M. J. Jutras, P. Fries, E. A. Buffalo, Oscillatory activity in the monkey hippocampus during visual exploration and memory formation. Proceedings of the National Academy of Sciences 110, 13144–13149 (2013).

13. H. S. Courellis, et al., Spatial encoding in primate hippocampus during free navigation. PLOS Biology 17, e3000546 (2019).

14. D. Mao, et al., Spatial modulation of hippocampal activity in freely moving macaques. Neuron 109, 3521 (2021).

15. S. Abbaspoor, A. T. Hussin, K. L. Hoffman, Theta- and gamma-band oscillatory uncoupling in the macaque hippocampus. eLife 12, e86548 (2023).

16. A. Jabès, P. B. Lavenex, D. G. Amaral, P. Lavenex, Postnatal development of the hippocampal formation: A stereological study in macaque monkeys. J. Comp. Neurol. 519, 1051–1070 (2011).

17. A. A. Liu, et al., A consensus statement on detection of hippocampal sharp wave ripples and differentiation from other fast oscillations. Nat Commun 13, 6000 (2022).

18. Y. Senzai, G. Buzsáki, Physiological Properties and Behavioral Correlates of Hippocampal Granule Cells and Mossy Cells. Neuron 93, 691–704.e5 (2017).

19. T. Donoghue, et al., Parameterizing neural power spectra into periodic and aperiodic components. Nat Neurosci 23, 1655–1665 (2020).

20. J. B. Caplan, J. R. Madsen, S. Raghavachari, M. J. Kahana, Distinct Patterns of Brain Oscillations Underlie Two Basic Parameters of Human Maze Learning. Journal of Neurophysiology 86, 368–380 (2001).

21. J. B. Caplan, et al., Human θ Oscillations Related to Sensorimotor Integration and Spatial Learning. J. Neurosci. 23, 4726 (2003).

22. A. M. Hughes, T. A. Whitten, J. B. Caplan, C. T. Dickson, BOSC: A better oscillation detection method, extracts both sustained and transient rhythms from rat hippocampal recordings. Hippocampus 22, 1417–1428 (2012).

23. L. L. Colgin, et al., Frequency of gamma oscillations routes flow of information in the hippocampus. Nature 462, 353–357 (2009).

24. S. Royer, et al., Control of timing, rate and bursts of hippocampal place cells by dendritic and somatic inhibition. Nat Neurosci 15, 769–775 (2012).

25. F. Cacucci, C. Lever, T. J. Wills, N. Burgess, J. O’Keefe, Theta-Modulated Place-by-Direction Cells in the Hippocampal Formation in the Rat. J. Neurosci. 24, 8265–8277 (2004).

26. J.-M. Alonso, L. M. Martinez, Functional connectivity between simple cells and complex cells in cat striate cortex. Nat Neurosci 1, 395–403 (1998).

27. K. Diba, A. Amarasingham, K. Mizuseki, G. Buzsáki, Millisecond Timescale Synchrony among Hippocampal Neurons. J. Neurosci. 34, 14984 (2014).

28. D. F. English, et al., Pyramidal cell-interneuron circuit architecture and dynamics in hippocampal networks. Neuron 96, 505–520.e7 (2017).

29. S. B. McHugh, et al., Offline hippocampal reactivation during dentate spikes supports flexible memory. Neuron 112, 3768–3781.e8 (2024).

30. A. Aljishi, N. E. Willhite, S. S. Cooper, K. L. Hoffman, Coordinated dynamics of Dentate Spikes and ripples in the hippocampus of sleeping macaques in 2025 Neuroscience Meeting Planner, Program No. PSTR1S2.0S, San Diego, CA, Society for Neuroscience., (2025).

31. W. E. Skaggs, et al., EEG Sharp Waves and Sparse Ensemble Unit Activity in the Macaque Hippocampus. Journal of Neurophysiology 98, 898–910 (2007).

32. T. K. Leonard, et al., Sharp Wave Ripples during Visual Exploration in the Primate Hippocampus. J. Neurosci. 35, 14771–14782 (2015).

33. J. S. Farrell, E. Hwaun, B. Dudok, I. Soltesz, Neural and behavioural state switching during hippocampal dentate spikes. Nature 628, 590–595 (2024).

34. M. Stewart, S. E. Fox, Hippocampal theta activity in monkeys. Brain Res 538, 59–63 (1991).

35. M. A. Belluscio, K. Mizuseki, R. Schmidt, R. Kempter, G. Buzsáki, Cross-Frequency Phase– Phase Coupling between Theta and Gamma Oscillations in the Hippocampus. J. Neurosci. 32, 423–435 (2012).

36. E. W. Schomburg, et al., Theta Phase Segregation of Input-Specific Gamma Patterns in Entorhinal-Hippocampal Networks. Neuron 84, 470–485 (2014).

37. A. B. L. Tort, et al., Dynamic cross-frequency couplings of local field potential oscillations in rat striatum and hippocampus during performance of a T-maze task. Proc. Natl. Acad. Sci. U.S.A. 105, 20517–20522 (2008).

38. L. L. Colgin, Theta–gamma coupling in the entorhinal–hippocampal system. Current Opinion in Neurobiology 31, 45–50 (2015).

39. C. J. McBain, A. Fisahn, Interneurons unbound. Nat Rev Neurosci 2, 11–23 (2001).

40. B. Rudy, G. Fishell, S. Lee, J. Hjerling-Leffler, Three groups of interneurons account for nearly 100% of neocortical GABAergic neurons. Developmental Neurobiology 71, 45–61 (2011).

41. A. Kepecs, G. Fishell, Interneuron cell types are fit to function. Nature 505, 318–326 (2014).

42. L. Roux, G. Buzsáki, Tasks for inhibitory interneurons in intact brain circuits. Neuropharmacology 88, 10–23 (2015).

43. Saleem, K.S., and Logothetis, N.K., A Combined MRI and Histology Atlas of the Rhesus Monkey Brain in Stereotaxic Coordinates, Second edition (Academic Press). (2012).

44. M. Pachitariu, S. Sridhar, J. Pennington, C. Stringer, Spike sorting with Kilosort4. Nat Methods 21, 914–921 (2024).

45. International Brain Laboratory, Spike sorting pipeline for the International Brain Laboratory. International Brain Laboratory (2022). Available at: https://figshare.com/articles/online_resource/Spike_sorting_pipeline_for_the_International_Brain_Laboratory/19705522/4.

46. E. K. Lee, et al., Non-linear dimensionality reduction on extracellular waveforms reveals cell type diversity in premotor cortex. eLife 10, e67490 (2021).

47. P. C. Petersen, J. H. Siegle, N. A. Steinmetz, S. Mahallati, G. Buzsáki, CellExplorer: A framework for visualizing and characterizing single neurons. Neuron 109, 3594–3608.e2 (2021).

48. S. Shinomoto, et al., Relating Neuronal Firing Patterns to Functional Differentiation of Cerebral Cortex. PLoS Comput Biol 5, e1000433 (2009).

49. R. Oostenveld, P. Fries, E. Maris, J.-M. Schoffelen, FieldTrip: Open source software for advanced analysis of MEG, EEG, and invasive electrophysiological data. Computational Intelligence and Neuroscience 2011 (2011).

50. P. P. Mitra, B. Pesaran, Analysis of dynamic brain imaging data. Biophysical Journal 76, 691– 708 (1999).

51. A. Grossmann, J. Morlet, “Decomposition of functions into wavelets of constant shape, and related transforms” in Mathematics + Physics, (WORLD SCIENTIFIC, 1985), pp. 135–165.

52. D. B. Percival, A. T. Walden, Spectral Analysis for Physical Applications: Multitaper and Conventional Univariate Techniques (Cambridge University Press, 1993).

53. K. H. Pettersen, A. Devor, I. Ulbert, A. M. Dale, G. T. Einevoll, Current-source density estimation based on inversion of electrostatic forward solution: Effects of finite extent of neuronal activity and conductivity discontinuities. Journal of Neuroscience Methods 154, 116–133 (2006).

54. R. M. M. Santiago, et al., Waveform-based classification of dentate spikes. Sci Rep 14, 2989 (2024).

55. D. Tingley, G. Buzsáki, Routing of Hippocampal Ripples to Subcortical Structures via the Lateral Septum. Neuron 105, 138–149.e5 (2020).

56. A. B. L. Tort, R. Komorowski, H. Eichenbaum, N. Kopell, Measuring phase-amplitude coupling between neuronal oscillations of different frequencies. J Neurophysiol 104, 1195–1210 (2010).

57. E. Maris, R. Oostenveld, Nonparametric statistical testing of EEG- and MEG-data. J Neurosci Methods 164, 177–190 (2007).

58. K. Mizuseki, K. Diba, E. Pastalkova, G. Buzsáki, Hippocampal CA1 pyramidal cells form functionally distinct sublayers. Nat Neurosci 14, 1174–1181 (2011).

59. E. Stark, et al., Pyramidal Cell-Interneuron Interactions Underlie Hippocampal Ripple Oscillations. Neuron 83, 467–480 (2014).

